# Biochemical and Molecular Characterization, and Bioprospecting of Drought Tolerant Actinomycetes from Maize Rhizosphere Soil

**DOI:** 10.1101/2020.05.13.094003

**Authors:** Chinenyenwa Fortune Chukwuneme, Olubukola Oluranti Babalola, Funso Raphael Kutu, Omena Bernard Ojuederie

## Abstract

Drought is a major limitation to maize cultivation around the globe. Seven actinomycetes strains were isolated from maize rhizosphere soils in Mahikeng, North-West Province, South Africa. The isolates were biochemically characterized and identified with 16S rRNA gene sequence analysis. Isolates were also screened *in vitro* for abiotic stress tolerance to different concentrations of NaCl, pH, and polyethylene glycol (PEG 8000), as well as for biosynthesis of drought tolerance genes namely Glutathione peroxidase (*GPX*), Glycine-rich RNA binding protein (*GRP*), Desiccation protectant protein (*DSP*), Guanosine triphosphate binding protein (*GTP*) and plant growth-promoting genes:1-aminocyclopropane-1-carboxylate deaminase (*accd*) and siderophore biosynthesis (*Sid*). About 71.43% of isolates were of the genus *Streptomyces* (99-100% similarity), while 14.29% belong to the genus *Arthrobacter* (R15) and 14.29% to the genus *Microbacterium* (S11) respectively (99% similarity). Five isolates had their optimum growth at 35°C. *Arthrobacter arilaitensis* (R15) grew and tolerated 5%, 10%, and 20% PEG at 120 h. Root length increased by 110.53% in PEG treated maize seeds (−0.30 MPa) inoculated with *Streptomyces pseudovenezuelae (*S20) compared to the un-inoculated control. Likewise, germination percentage and vigor index increased by 37.53% and 194.81% respectively in PEG treated seeds inoculated with S20 than the un-inoculated PEG treated seeds. ACC deaminase gene was amplified in all the isolates, while the gene for siderophore biosynthesis was amplified in 85.71% of the isolates. Genes for the synthesis of GPX, GRP, DSP and GTP were amplified in *Arthrobacter arilaitensis* (R15) and *Streptomyces pseudovenezuelae* (S20) which lacked GTP. The amplification of drought-tolerant and plant growth-promoting primers indicates the possible presence of these genes in the isolates. These isolates have the potential for use as bio-inoculants, not only to improve drought tolerance in maize but also to be utilized as biofertilizers and biocontrol agents to facilitate growth promotion.

## Introduction

As the world’s climatic conditions change because of massive increases in the world’s population and global industrialization, more agricultural land is being lost to drought. Loss of arable land due to drought is a problem that is becoming common in many regions of the world as it is expected to result in a 30% loss of land by 2021 and greater than 50% by 2050 [1, 2]. Current studies have shown that it is not only limited to arid areas but also occurs in temperate countries [3, 4]. Plants are constantly subjected to different abiotic and biotic stresses such as drought, salinity, flooding, high temperatures, toxic metals, radiation, insect, fungi, bacteria, and viruses [5, 6]. Drought, a prominent abiotic stress-reducing plant growth and crop yield, results in a substantial decrease in agricultural productivity. Drought causes considerable loss of crops worldwide, bringing about more than 50% reduction in the average yields of most major food crops, and is therefore a major threat to global food security [2]. Plants respond to the deleterious effects of drought stress by synthesizing various defense-related proteins, and by modifying their physiological and metabolic activities [7, 8].

To eradicate the problem of water scarcity, modern agro-biotechnological strategies are being implemented to improve drought tolerance in plants. These technologies include genetic engineering, germplasm screening, plant breeding, etc., which have resulted in the development of hybrid products that are widely grown across different parts of the globe [9-11]. Nevertheless, abiotic stress tolerance mechanisms are complex, making the task of introducing new tolerant varieties tough and strenuous [10]. Other agricultural methods to reduce the negative effects caused by drought are conservation tillage, soil amelioration, and mulching, but these methods are not only difficult but also time-consuming [12]. As agricultural activities spread to less fertile and marginal soils, more attention is being channeled to interventions that are capable of increasing water use efficiency in plants through biotechnology and improved agronomic practices to satisfy the increasing demand for food [13, 14].

Certain soil microorganisms known as plant growth-promoting bacteria (PGPB) can aid plants to overcome the drastic impact of different ecological stresses [4, 15-17]. This group of bacteria enhances plant growth directly by providing them with phytohormones, namely: gibberellins, cytokinins, and indole-acetic acid; and aiding in the acquisition of nutrients such as phosphorus and atmospheric nitrogen [18]. They indirectly promote plant growth by reducing plant ethylene levels by the activity of 1-aminocyclopropane-1-carboxylate (ACC) deaminase enzyme [18-20]. Likewise, they can act as biocontrol agents by preventing or reducing the damage caused by fungi or bacteria on plants [12, 17, 18, 21, 22]. Therefore, an understanding and improvement of plant growth and productivity under restricted water availability using PGPB is crucial.

Bacteria have evolved several mechanisms to tackle the damages caused by drought on plants. These mechanisms may include modifications of phytohormones which play a major role in helping plants escape or survive abiotic stresses [23, 24], alteration in plant root morphology, accumulation of osmolytes, alteration of plant antioxidant defense mechanisms, production of exopolysaccharides and the identification of candidate genes responsible for plant growth promotion and drought tolerance [8, 25]. Bacteria possessing these traits can be isolated and used to improve plant growth under stress conditions. For a successful application of this strategy, a good knowledge of the ability of these organisms to tolerate drought is required. The successful isolation of bacteria for drought tolerance in plants has been proven and reported from several types of research, but very few on the genus actinomycetes. Moreover, only a few of these studies reported on the molecular mechanisms of these bacteria in drought tolerance. Identifying the genetic make-up of these bacteria as well as the evidence of genes encoding drought tolerance may be of help in understanding the mechanism of bacteria-mediated tolerance to abiotic stress.

Information on the actinomycetes strains obtained from this study will go a long way in the selection of beneficial actinomycetes strains that can be used for plant growth enhancement under drought stress. The study aimed to isolate and characterize actinomycetes from maize rhizosphere soil and determine their drought tolerance abilities. We also report the design of primers capable of amplification of specific genes encoding proteins involved in drought tolerance as well as the specific genes encoding proteins responsible for plant growth promotion and drought tolerance.

## Materials and methods

### Collection of soil samples

Four (4) rhizosphere soil samples were collected from two different maize fields, (i) behind Animal Health Center, North-West University, Mafikeng Campus, and (ii) North-West University Agricultural Farm, Molelwane, South Africa [26]. Soil samples were collected by carefully uprooting dry maize plants and shaking the plants to remove soils loosely adhered to the plant roots. Soils tightly adhered to the roots were aseptically collected in sterile plastic bags and transported to the Microbial Biotechnology Laboratory in a cooler box. Collected samples were stored at -20°C for further analysis.

### Isolation and selection of bacteria

Bacterial isolation was carried out by suspending 1 g of each soil sample separately in 9 ml of sterile saline solution (0.85%) in 20 ml sterile test-tubes. The test-tubes were thoroughly shaken using a vortex machine and serial dilution method used to isolate the actinomycetes strains from the soil samples [27, 28]. Aliquots of 200 µl from each dilution were spread using a glass rod on Actinomycete Isolation Agar (AIA) plates (pH, 7.2) supplemented with cycloheximide (20 mg/l) and nalidixic acid (100 mg/l) to minimize fungal and bacterial growths respectively (these were performed in triplicate). Plates were incubated at 25°C for 10 days for optimum growth. Isolated bacteria with varying color and shape were randomly selected and repeatedly streaked on freshly prepared Yeast extract-malt extract agar (ISP-medium 2) plates to obtain pure actinomycetes cultures. Pure cultured bacterial strains were maintained on agar slants at 4°C. Seven pure actinomycetes isolates were obtained which were further characterized.

### Morphological characterization

The morphological and cultural analysis of rhizospheric actinomycetes were performed according to the standard of the International Streptomyces Project (ISP) [29, 30]. Isolates were identified up to the genus level by checking the color of the aerial and substrate mycelium and other traits such as the color of colonies on Petri plates and spore mass color [31]. Biochemical tests were carried out on nitrate reduction, starch hydrolysis, catalase activity, and utilization of carbohydrate sources. Isolates were further characterized by following the keys of Bergey’s Manual of Determinative Bacteriology [32].

### Biochemical characterization

#### Nitrate reduction test

A loopful of spores from selected bacterial isolates was inoculated into 20 ml test-tubes containing 5 ml of sterilized nitrate broth and incubated at 25°C for 8 days. Controls were test-tubes containing broth without inoculation. On the 8^th^ day of the incubation period, broths were tested for the presence of nitrate by the addition of two drops of reagent A (α-naphthylamine solution) and two drops of reagent B (sulfanilic acid solution) into 1 ml of broth or 1 ml of control. A change in color of the broth to pink, orange, or red indicated the presence of nitrate. Results were confirmed by the addition of a pinch of zinc dust after the reagents were added. A change in color from red to the original color of the broth confirms the result.

#### Utilization of carbohydrate sources

The seven selected actinomycetes isolates were tested for their ability to utilize various carbon compounds as energy sources using ISP-medium 9 as recommended by Shirling and Gottlieb [29]. The carbon sources used: D-galactose, D-glucose, D-xylose, sucrose, D-mannitol, lactose, and fructose, were filter-sterilized using a micro-membrane filter (20 µm/cm) to ensure they were free from various contaminants. All inoculated tubes were incubated at 25°C for 8 days. Acid production by the isolates from the above carbon sources was also studied. The positive utilization of a carbon source was considered when the growth of an isolate on a tested carbon source was significantly better than the growth on the basal medium without a carbon source [33], while growth similar or less than growth on basal medium without a carbon source was considered a negative utilization [28].

#### Catalase production test

Luria-Bertani (LB) agar slants were inoculated with each bacterial isolate. An un-inoculated LB slant was used as the control. All tubes were incubated at 30°C for 5 days. On the 5^th^ day, a single colony from each inoculated tube was placed on a sterile glass slide and held at an angle while 2-3 drops of hydrogen peroxide (H_2_O_2_) which flowed over the growth of each culture on the glass slide. Catalase production was indicated by the production of oxygen bubbles within one minute after the addition of H_2_O_2_.

#### Test for hydrolysis of starch

A loopful of each bacterial isolate was streaked on starch agar plates [34] in triplicate; plates were incubated at 30°C for 8 days. On the 8^th^ day, the iodine solution was added to each culture plate. After about a minute, excess iodine was carefully poured out from the culture plates. Controls consisted of un-inoculated starch agar plates. A yellow zone around the colony in a dark blue medium was considered positive for starch hydrolysis, while the absence of a yellow zone was considered negative.

#### Test for casein hydrolysis

The ability of selected isolates to hydrolyze casein was determined by spot-inoculating prepared skimmed milk agar plates with a loopful of each bacterial isolate. Un-inoculated skimmed milk agar plates served as the control for this experiment, and all experiments were replicated thrice. All plates (inoculated and control) were incubated at 30°C for 5-8 days. Observation of a clear zone around the bacterial growth on the agar culture plates indicated evidence of casein hydrolysis.

#### Molecular characterization

##### Genomic DNA extraction, quantification, and PCR amplification of 16S rRNA gene

Bacterial isolates were grown in 20 ml of ISP-1 medium in 50 ml Eppendorf tubes containing 5% PEG 8000 with constant agitation (150 rpm) at a temperature of 25°C for 7 days for optimum growth. Bacteria were harvested by centrifuging at maximum speed for 10 min. Supernatants were discarded, and cells were collected and re-suspended in sterile distilled water. The total DNA from each bacterium was extracted using a DNA extraction kit (ZR soil microbe DNA MiniPrep™ kit (Zymo Research, USA) according to the manufacturer’s instructions [35]. DNA purity and yield (ng/µl) was determined using a NanoDrop Spectrophotometer (Thermo Scientific, DE, USA).

Actinomycetes identity of the 7 isolates was confirmed by amplification of the 16S rRNA gene using the universal 16S rRNA primers for bacteria fD1, rP2 [28, 36] in a thermal cycler (BioRad). Details of primers are shown in Supplementary Table S1. Amplifications were performed in a final volume of 25 µl consisting of 12.5 µl of 2x PCR Master Mix (0.05 U/µl *Taq* DNA polymerase, 4 mM MgCl_2_ and 0.4 mM dNTPs (Fermentas)), 1 µl of genomic DNA template, 0.5 µl of each of the forward and reverse primers and 10.5 µl of nuclease-free water [35]. The PCR cocktail was subjected to 30 cycles in a C1000 thermal cycler (BioRad) with an initial denaturation temperature at 95°C for 3 min; denaturation at 96°C for 45 s; annealing temperature at 56°C for 30 s; extension at 72°C for 2 min and a final extension at 72°C for 5 min.

### Purification of PCR product, sequencing, and phylogenetic analysis

Purification and sequencing of PCR products were performed by Inqaba Biotechnical Industrial (Pty) Ltd, Pretoria, South Africa using PRISM™ Ready Reaction Dye Terminator Cycle Sequencing Kit [37]. The analysis of sequences and construction of the phylogenetic tree was performed according to the methods described by Aremu and Babalola [37]. The chromatograms obtained from the sequencing reaction were analyzed for good quality sequences using the Chromas Lite version 6.5 [38]. The chromatograms obtained were edited with Bio Edit Sequence Alignment Editor [39], and consensus sequences were generated. Blast search was done on the consensus sequences obtained in the NCBI database (www.ncbi.nlm.nih.gov) using the Basic Alignment Search Tool (BLASTn) for homology for identification of the bacteria [37]. Sequences obtained were deposited in GenBank.

The evolutionary history was deduced by using the Maximum Likelihood approach based on the Tamura-Nei model [40]. The tree with the highest log likelihood (−6740.13) was obtained. The percentage of trees in which the related taxa grouped together appears next to the branches. Initial tree(s) for the heuristic search were obtained automatically by applying Neighbor-Joining and BioNJ algorithms to a matrix of pairwise distances estimated using the Maximum Composite Likelihood (MCL) approach, with the topology with superior log likelihood value selected. A total of 36 nucleotide sequences were used for the analysis, which included 1st+2nd+3rd+Noncoding. Gaps and missing data were removed from all positions. The final dataset contained a total of 530 positions. Evolutionary analyses were performed in MEGA7 [41].

### PCR amplification of specific genes encoding proteins involved in drought tolerance and plant growth promotion (PGP) in bacterial isolates

#### Design and development of specific primers encoding drought tolerance genes

Specific primers encoding drought tolerance genes used in this study were designed according to the protocol of Aremu and Babalola [37]. Database searching was through the National Center for Biotechnology Information (NCBI) at http://www.ncbi.nlm.nih.gov/ [37]. Retrieved sequences were glutathione peroxidase, Desiccation protectant protein, guanosine triphosphate binding protein, and glycine-rich RNA binding protein. The nucleotide sequences of these genes encoding for drought tolerance from NCBI were saved in FASTA format and copied into the Bio Edit Sequence Alignment Editor (Version 7.0.9.0) [39]. ClustaI W 2.0 algorithm [42] was used to perform Multiple Sequence alignments (MSA). Stringency was varied to achieve as few gaps and mismatches as possible. Many regions with an exceedingly high degree of sequence similarity were obtained after an alteration of the stringency. The consolidation of MSA was performed following the method of Aremu and Babalola [37]. Subsequently, consolidated trials were aligned with each other, and low similarity sequences discarded. In order to design primers for drought tolerance from the highly conserved regions, the files were opened in Bio Edit and Primers designed by means of Primer3Plus interface (http://frodo.wi.mit.edu/) and the most appropriate primers chosen using the standards for good primer design [37, 43]. The chosen primer sequences were tested for potential hairpins structures, cross homology, cross dimer, and self-dimer. These were further tested using Gene Infinity Platform for binding similarities to the priming sites (delta G values), as described by Aremu and Babalola [37]. *In silico* PCR in Gene Infinity Platform was employed to determine the specificity of the primers [37], and the suitability of the primers to target drought tolerance encoding genes checked using NCBI Blast.

#### PCR amplification of drought tolerance genes by specific primers

Primers were assessed for varying annealing temperatures using a gradient PCR machine from 45 to 60°C. Reaction volumes were 25 µl which consisted of 12.5 µl of 2x PCR Master-Mix (0.05 U/µl *Taq* DNA polymerase, 4 mM MgCl_2_ and 0.4 mM dNTPs (Fermentas) [35], 1 µl of genomic DNA template (10 ng/ reaction mixture), 0.5 µl of each of the forward and reverse primers and 10.5 µl of nuclease-free water [35]. The PCR program began with a 95°C hot start for 10 min, followed by denaturation at 94°C for 30 s/ cycle, a 45-60°C annealing temperature for 30 s, 72°C elongation step for 1 min, a final elongation step of 72°C for 10 min and a holding period at 4°C for infinity.

#### PCR amplification of plant growth-promoting genes

The actinomycetes isolates were screened for amplification of the Siderophore (*Sid*) and ACC deaminase (*accd*) genes associated with plant growth promotion. The PCR for siderophore and ACC deaminase genes were performed as follows: 2 µl (about 10 ng) of each DNA extract was amplified with 12.5 µl of 2x PCR Master Mix (0.05 U of *Taq* DNA polymerase4 mM MgCl_2_ and 0.4 mM dNTPs (Fermentas), 1 µM of each primer and 8.5 µl of nuclease-free water, in a 25 µl reaction mixture using a C1000 thermal cycler (BioRad) with the following PCR programme: 30 cycles of denaturation at 94°C for 1 min, annealing temperatures of 55° (*sid*) 52°C (*accd*) for 45 s, an extension at 72°C for 2 min and a final extension step at 72°C for 7 min. Supplementary Table S1 shows the details of primers used for the amplification of 16S rRNA (F1r2), and plant growth-promoting genes (*accd* and *Sid*) used in the present study.

#### Agarose gel electrophoresis

The quality of genomic DNA and the PCR products were checked on 1% (w/v) agarose gel made by dissolving 1.5 g of agarose (Bio-Rad, SA) in 150 ml of 1X Tris-acetate-ethylenediaminetetraacetate (TAE, pH 8). The mixture was placed in a microwave and heated for 3 min and left to cool. Subsequently, 10 µl of Ethidium bromide (EtBr) was added to the molten gel and then poured in a gel casting tray with combs and allowed to solidify [35]. The combs were removed after solidification, and the gel carefully placed in the electrophoresis tank containing 1X TAE buffer (40 nM Tris, 20 mM Acetic acid and 100 mM EDTA, pH 8.0) [35]. DNA samples were prepared by mixing 7 µl of DNA template, and 3 µl of 6x DNA loading dye (Fermentas), 5 µl PCR products and 2 µl DNA ladders (1 kb or 100 bp) were also taken. Samples were carefully loaded into the preformed wells in the gel, and the samples run for 60 min at 80 V. Results were visualized and photographed using a ChemDoc™ MP System (Bio-Rad Laboratories, Hercules, CA, USA) [37].

#### Effect of temperature on the growth of bacteria

Temperature effect on bacterial growth was done by growing 10 µl (3.8 x 10^6^) of each bacterial isolate in 10 ml of sterilized ISP-1 medium containing 5% PEG-8000 and incubated at different temperatures (25°C, 30°C, 35°C, 40°C) under shaking conditions (150 rpm) for 7 days [26]. The OD of each bacterial isolate was measured at 600 nm using a UV Spectrophotometer (Merck, SA) [35].

#### Effect of pH on the growth of bacteria

The effect of pH on bacterial growth was determined by growing 10 µl of each bacterial culture in test tubes containing 10 ml of sterilized ISP-1 medium supplemented with 5% PEG 8000. The pH of the media was adjusted to 3, 5, 7, 9, and 11 using 1 N HCl and 1 N NaOH before autoclaving. Inoculated tubes were incubated at 25°C under shaking conditions (150 rpm) for 7 days [26], after which the OD of each culture was measured using a UV spectrophotometer (Thermo Spectronic, Merck, SA) [35].

#### Effect of NaCl on the growth of bacteria isolates

Tolerance to NaCl by bacteria was assessed according to the method of Ndeddy Aka and Babalola [35] by inoculating 20 µl of each bacterial isolate in 20 ml sterilized ISP-1 medium containing varying concentrations of NaCl (0.2, 0.4, 0.6, 0.8, and 1.0%). Inoculated tubes were incubated at 25°C for 7 days [26], and the OD was measured at 600 nm using a UV spectrophotometer (Thermo Spectronic, Merck chemicals, SA) [35]. An OD_600_ value greater than 0.1 is considered as a better growth for each bacterial isolate.

#### Drought tolerance abilities of bacterial isolates

To study the drought tolerance capacities of bacterial isolates used in this study, a 5 ml (3.8 x 10^6^ CFU ml^-1^) aliquot of freshly prepared cultures were inoculated into 150 ml cotton plugged flasks containing 50 ml sterilized ISP-1 medium supplemented with different concentrations (5, 10, 15 and 20%) of PEG 8000. The pH of the medium was adjusted to 7.2 before autoclaving at 121°C for 15 min. Inoculated flasks were incubated at room temperature with constant shaking (150 rpm) at different time intervals (24, 48, 72, 96, and 120 h). The growth of each bacterial isolate was obtained by measuring the OD at 600 nm using a UV Spectrophotometer (Thermo Spectronic, Merck chemicals, SA). The control for this experiment consisted of inoculated broth without PEG 8000, while un-inoculated broth served as blank. The experiment was performed in triplicate, and pooled data were statistically analyzed.

#### *In-vitro* seed germination assay of bacterial isolates

The drought tolerance abilities of four bacterial isolates S20, S12, R11, and S11 were tested using maize seeds of variety S0/8/W/ I137TNW//CML550 obtained from the Agricultural Research Council (ARC), South Africa [26]. Seeds were prepared following the method of Chukwuneme, Babalola (26). Afterward, 10 clean Petri plates in triplicates were prepared by placing two filter papers at the bottom of each plate. The first 5 plates contained 10 ml of sterile tap water, while the second 5 plates contained 10 ml of -0.30 MPa PEG 8000 solution. For the first 5 plates, sterile seeds (10) previously immersed in 20 ml (4.0 x 10^7^ CFU ml^-1^) of each bacterial suspension and shaken in a rotary shaker for 4 h at 150 rpm were placed in 4 plates out of the first 5 plates. The remaining plate in this category contained maize seeds previously immersed in distilled water and severed as the control for the experiment. For the second set of 5 plates, 10 sterile seeds previously immersed in 20 ml of (4.0 x 10^7^ CFU/ml) of each bacterial suspension and shaken in a rotary shaker for 4 h at 150 rpm were placed in 4 plates. In contrast, the remaining plate in this category contained 10 seeds without bacterial inoculation distilled water were placed in each plate for 4 h. 10 ml of sterile distilled water or 10 ml of -0.30 MPa PEG 8000 solution as the case may be was added to the plates every 2 days, until the end of the experiment. The following codes were used for the first 5 plates: Plate 1: S20+H_2_0, plate 2: S11+H_2_O, plate 3: R11+H_2_0, plate 4: S11+H_2_0, and plate 5: M+H_2_O. The codes for the second 5 plates containing PEG 8000 were as follows: Plate 6: S20+PEG, plate 7: S11+PEG, plate 8: R11+PEG, plate 9: S11+PEG, and plate 10: M+PEG. Seeds were left to grow for 12 days after which 5 best seedlings were selected from each plate for measurement of growth parameters.

The experiment was performed in triplicate. The percentage germination and vigor index of seedlings were estimated as previously described by Chukwuneme, Babalola [26] as follows:

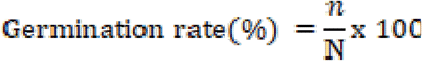

Where n is the number of germinated seeds after 10 days, and N is the total number of seeds [26].

Vigor index = % germination x total length of seedling (shoot length + root length) [26]

### Data analysis

All quantitative data obtained from this study were analyzed by one-way analysis of variance (ANOVA) using the Statistical Analysis Software (SAS), Version 9.4 [26, 44]. Significant mean differences were compared using New Duncan’s Multiple Range Test [26].

## Results

### Isolation and characterization of actinomycetes isolates

#### Morphological and biochemical traits

Seven actinomycetes were isolated from maize rhizosphere soils and characterized based on the morphology of the colonies, their ability to form aerial hyphae, and substrate mycelium as well as their biochemical properties. Most isolates showed a slow growth rate except for isolate R15, which showed moderate growth on the medium. After 10 days’ incubation period, colonies were observed to be white, brownish-white, and gray (**Table 1**). The biochemical properties disclosed that all tested bacterial isolates had a positive reaction for catalase activity, nitrate reduction, starch hydrolysis, and the utilization of glucose as a carbon source. Moreover, only two tested bacterial isolates (S4 and S7) hydrolyzed casein. From our results, four isolates (S12, S4, S11, and S20) utilized D-galactose, four (R15, S7, S4, and R11) utilized the D-xylose, five (R15, S12, S4, S11, and S20) utilized sucrose and two (S4 and R11) utilized D-mannitol. The results obtained also showed that all the isolates used fructose except isolate S7, while this isolate was the only one that utilized lactose among all tested bacterial isolates (**Table 2**).

**Table 1:** Morphological properties of isolated rhizospheric actinomycetes.

| Isolates | Growth and colony nature | Color of aerial mycelium | Color of substrate mycelium | Formation of pigments |
| --- | --- | --- | --- | --- |
| S4 | Slow and smooth | gray | Brown | No |
| S7 | Slow and smooth | gray | Light brown | No |
| S11 | Slow and firm | White | Yellowish-brown | No |
| S12 | Slow and firm | White | Light brown | No |
| S20 | Slow and firm | White | Yellowish-brown | Yes |
| R11 | Slow and firm | gray | Light brown | No |
| R15 | Moderate and powdery | Brownish white | White | Yes |

**Table 2.** Biochemical and physiological features of bacterial isolates.

| Biochemical and physiological properties | Bacterial isolates |  |  |  |  |  |  |
| --- | --- | --- | --- | --- | --- | --- | --- |
|  | S4 | S7 | S11 | S12 | S20 | R11 | R15 |
| Catalase activity | + | + | + | + | + | + | + |
| Nitrate reduction | + | + | + | + | + | + | + |
| Casein hydrolysis | + | - | + | - | - | - | - |
| Starch hydrolysis | + | + | + | + | + | + | + |
| **Utilization of carbohydrates** |  |  |  |  |  |  |  |
| D-galactose | + | - | + | + | + | - | - |
| D-glucose | + | + | + | + | + | + | + |
| D-xylose | + | + | - | - | - | + | + |
| D-mannitol | + | - | - | - | - | + | - |
| Sucrose | + | - | + | + | + | - | + |
| Lactose | - | + | - | - | - | - | - |

Note: S and R stand for bacterial isolates

#### Phylogenetic analysis of actinomycetes isolates

The 7 actinomycetes isolates were identified using 16S rRNA gene sequencing and BLASTn analysis (**Table 3**). The BLAST search inferred that the isolates belong to the GC-rich Actinomycetales. The BLASTn homology result revealed that the 16S rRNA gene sequences of 5 isolates among the 7 isolates were confirmed as *Streptomyces* spp. Isolates R15 and S11 were confirmed as *Arthrobacter* and *Microbacterium*, respectively. Isolates S7 had a 100% similarity with *Streptomyces luteogriseus* while isolate S4 had 99.9% similarity with *Streptomyces werraensis.* Isolates R11, R15, S11, and S12 showed 99% similarity to *Streptomyces indiaensis, Arthrobacter arilaitensis, Microbacterium oxydans*, and *Streptomyces* spp. respectively, while S20 had 93% similarity to *Streptomyces pseudovenezuelae* (**Table 3**). The gene sequences of all isolates were submitted to GenBank (NCBI) with the following accession numbers: MG547867 (S4), MG669347 (S7), MG547868 (R11), MG547869 (R15), MG547870 (S20), MG547870 (S11) and MG640369 (S12) (**Table 3**).

**Table 3:** Partial 16S rRNA sequence identification of the species closely associated with the actinomycetes isolates using BlastN program

| Isolates | Accession numbers | Query cover | E-value | Sequence similarity (%) | Blast ID (closest representative species) |
| --- | --- | --- | --- | --- | --- |
| S4 | MG547867 | 100 | 0 | 99.9 | *Streptomyces werraensis* |
| S7 | MG669347 | 100 | 0 | 100.0 | *Streptomyces luteogriseus* |
| S11 | MG640368 | 100 | 0 | 99.7 | *Microbacterium oxydans* |
| S12 | MG640369 | 100 | 0 | 99.6 | *Streptomyces* sp. |
| S20 | MG547870 | 100 | 0 | 99.1 | *Streptomyces pseudovenezuelae* |
| R11 | MG547868 | 100 | 0 | 99.1 | *Streptomyces indiaensis* |
| R15 | MG547869 | 99 | 0 | 99.5 | *Arthrobacter arilaitensis* |
| Out-group | MN685265 | 97 | 0 | 78.9 | *Pseudomonas gessardii* |

Phylogenetic analyses using the Maximum Likelihood technique were computed with the 16S rRNA gene sequences of the 7 isolates (**Fig 1**). The phylogenetic tree revealed the relationship of the bacterial isolates with the closest reference bacteria species. The 16S rRNA sequences of the seven isolates were aligned to twenty-eight (28) reference sequences of the 16S rRNA gene of closely related taxa as retrieved from GenBank data library, and *Pseudomonas gessardii* strain OBE3 (MN685265) was used as the out-group (**Fig 1)**. The 7 actinomycetes isolates all belong to the phylum Actinobacteria, but 71.43% were of the *Actinomycetaceae* family and genus *Streptomyces*, while the other two isolates S11 and R15 were of the *Microbacteriaceae* and *Micrococcaceae* families and genera *Microbacterium* and A*rthrobacter*, respectively. All the isolates were assembled with their corresponding genus in the tree with high bootstrap values ranging from 94% to 100%. Isolate R11 with a 94% bootstrap value was closely related to *Streptomyces indiaensis* (FJ951435) while isolate R15 had remarkably close ancestry with *Arthrobacter arilaitensis* (JQ071516) and *Glutamicibacter arilaitensis* (MG547869 and MH130322). Isolate S4 was very distinct, with 100% bootstrap value and was closely related to *Streptomyces werraensis* (MH130305 and MK825539). Isolate S7 had a close association to three species of the genus *Streptomyces* with a 99% bootstrap value (*Streptomyces luteogriseus* (MH259082), *Streptomyces paradoxus* (MT072136) and *Streptomyces purpurascens* (MT084590) respectively. Isolate S12 showed high relatedness (98%) to *Streptomyces* sp. (MT255053), *Streptomyces parvulus* (MN901807), and Streptomyces rochei (MT255666). In comparison, isolates S11 and S20 had 97% bootstrap values revealing very close association to *Microbacterium oxydans* (LN8090040) and *Brachybacterium nesterenkovii* (KY979106) and *Streptomyces pseudovenezuale* (MF417392) and *Streptomyces griseus* (LT221172), respectively. PCR amplification of the 16S rRNA genes was successfully performed using F1r2 universal bacteria primers (**Supplementary Table S1**). 1 Kb DNA marker (GeneRuler™ DNA Ladder) was used to determine the band sizes (**Fig.2**).

**Fig 1.**
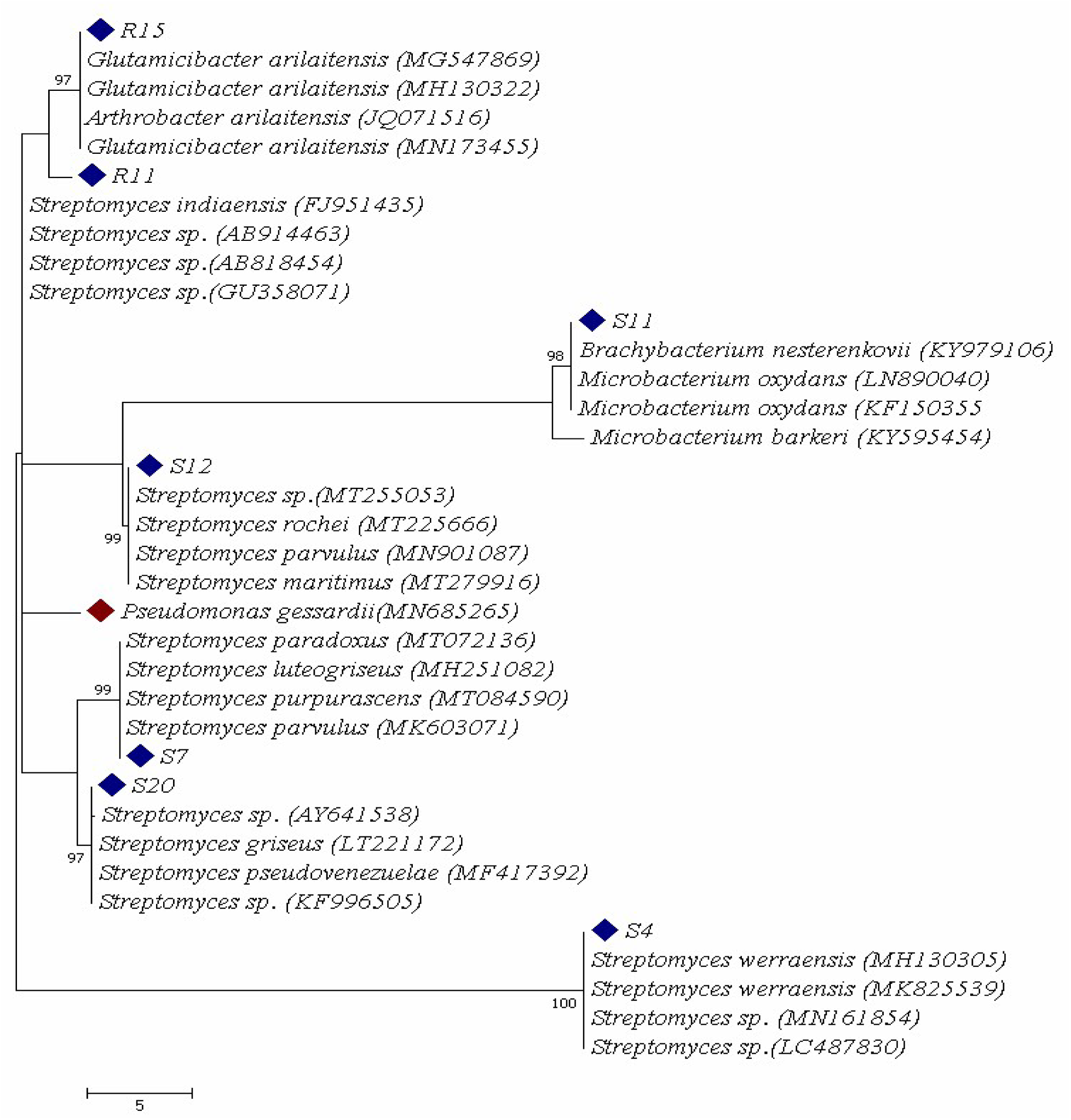
Maximum Likelihood tree of the 7 actinomycetes isolates and representative bacterial species of based on partial 16S rRNA gene sequences. Numbers at the nodes indicate the levels of bootstrap support based on 1000 resampled data sets. Only values greater than 50% are shown. The scale bar indicates 5 substitutions per nucleotide position. The isolates are indicated by a diamond shape

**Fig 2.**
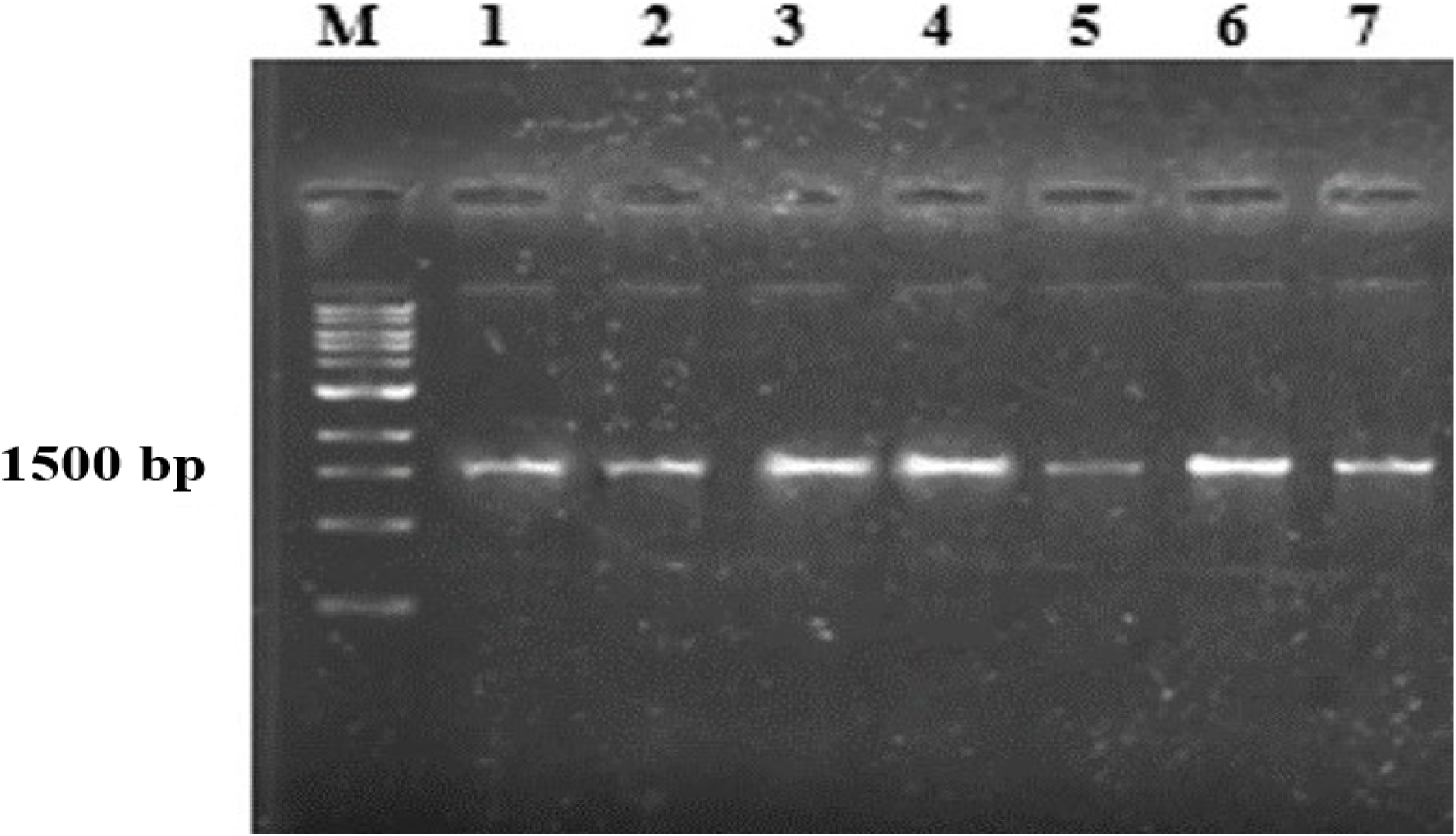
Amplified rRNA sequences of actinomycetes isolates using universal primer F1r2. Lane M= 1Kb DNA marker, Lane 1: S4, Lane 2: S7, Lane 3: S20, Lane 4: R11, Lane 5: R15, Lane 6: S12, and Lane 7: S11.

#### Primer design and amplification of drought-tolerant and PGP genes

The sets of primers used in this study (**Supplementary Table S2**), were developed around the species of bacteria that will be able to enhance abiotic stress tolerances in plants for their use in amplifying DNA in bacteria. Four primers from 16S sequences of bacteria were developed successfully. The designed primers were tested for binding affinities to the priming sites (delta G values) in the Gene Infinity Platform [37]. Tests showed that the designed primers as presented in supplementary Table S2 did not have cross-dimer, self-dimer, cross homology, and potential hairpin structures. All forward primers had appropriate G/C clamp at the 3’ ends, moderate melting temperatures, and their locations were past the 5’ end of the coding sequence [37]. The lengths of the primer sequences were moderate, assisting in precise binding to the target genes. The excellent specificity of the primers in the Gene Infinity Platform were confirmed by *in silico* PCR. Also, primer specificity performed in the NCBI Primer-BLAST gave the target drought tolerance genes. The PCR amplification of the *GPX* gene yielded the expected band size of 215 bp for isolates R15, S20, S7, and R11 while isolates S11, S12 and S4 yielded no amplification. (**Supplementary Fig S1A**). In the case of GRP (**Supplementary Fig S1B**), isolates R11, S20, S12, and R15 yielded the expected band size of 220 bp while isolates S4, S7, and S11 did not amplify. Isolates S20 and R15 amplified the *DSP* gene at the expected product size of 920 bp while no amplification observed for the other isolates at this size and the GTP gene was only amplified in isolate R15 at the expected band size of 668 bp (**Supplementary Fig. S1C**).

The ACC deaminase (*accd*) gene was amplified in all seven isolates (**Supplementary Fig S1A**) with the reference primers, as shown in Supplementary Table S1. The amplification of siderophore (*Sid*) gene (**Supplementary Fig S1B**) for the seven isolates was performed by PCR using reference primers (Supplementary Table S1). It was observed that isolates S20, R11, S12, R15, S11 and S4 amplified this gene. However, no amplification was observed for isolate S7 for this gene.

#### Effect of temperature on bacterial growth

The activity and growth of bacteria in the soil depend on the soil temperature, which affects cellular enzymes. In the present study, seven actinomycetes isolates were grown under different temperature regimes. The results obtained for each of the isolates at various temperatures are presented in Table 5. Bacterial growths were determined by measuring the optical density (OD) of each bacterium at 600 nm. All tested bacterial isolates grew at various temperatures, although there were variations in the growth pattern of each isolate. Isolate S4 had optimum growth at 35°C (0.65 ± 0.11) and its least growth at 25°C (0.29 ± 0.13) which was significantly lower than the growth recorded for other isolates at 25°C with optimum growth recorded for isolate S11 (0.68 ± 0.12). At 30°C, optimum bacterial growth was recorded for isolates S20 (0.56 ± 0.32), S12 (0.55 ± 0.16), and R15 (0.53 ± 0.19) which were not significantly different from one another but significantly higher than the growth obtained for other isolates with the least growth recorded in isolate S4 (0.34 ± 0.19). At a temperature of 35°C, isolate R15 had its optimum growth rate (0.91 ± 0.04) which was significantly (p < 0.001) higher than the growth recorded for other isolates, followed closely by isolate S12 (0.89 ± 0.15) with S20 having the least growth (0.27 ± 0.12). At this temperature (35°C), isolates R11 and S4 had their optimum growth rates (0.64 ± 0.12) and (0.65 ± 0.11), respectively. Most isolates had their optimum growth at 35°C except isolates S11 and S20 (**Table 5**). At the highest temperature of 40°C, a significant decrease in growth was observed for most isolates (R15, S11, S20, and S7). Nonetheless, isolate R11 had the highest growth (0.42 ± 0.11) followed by isolates S12 (0.40 ± 0.11) and S4 (0.32 ± 0.04).

**Table 5.** Effect of temperature on the growth of bacterial isolates.

| Isolates | Percentage growth at different temperatures (°C) |  |  |  |
| --- | --- | --- | --- | --- |
|  | 25 | 30 | 35 | 40 |
| S4 | 0.29 ± 0.13<sup>d</sup> | 0.34 ± 0.19<sup>b</sup> | 0.65 ± 0.11<sup>c</sup> | 0.32 ± 0.04<sup>abc</sup> |
| S7 | 0.39 ± 0.06<sup>cd</sup> | 0.47 ± 0.09<sup>ab</sup> | 0.81 ± 0.08<sup>b</sup> | 0.28 ± 0.02<sup>abc</sup> |
| S11 | 0.68 ± 0.12<sup>a</sup> | 0.47 ± 0.16<sup>ab</sup> | 0.46 ± 0.08<sup>d</sup> | 0.22 ± 0.02<sup>c</sup> |
| S12 | 0.61 ± 0.12<sup>ab</sup> | 0.55 ± 0.16<sup>a</sup> | 0.89 ± 0.15<sup>ab</sup> | 0.40 ± 0.11<sup>ab</sup> |
| S20 | 0.41 ± 0.19<sup>cd</sup> | 0.56 ± 0.32<sup>a</sup> | 0.27 ± 0.12<sup>e</sup> | 0.25 ± 0.13<sup>bc</sup> |
| R11 | 0.53 ± 0.18<sup>abc</sup> | 0.41 ± 0.25<sup>ab</sup> | 0.64 ± 0.12<sup>c</sup> | 0.42 ± 0.11<sup>a</sup> |
| R15 | 0.45 ± 0.19<sup>bcd</sup> | 0.53 ± 0.19<sup>a</sup> | 0.91 ± 0.04<sup>a</sup> | 0.21 ± 0.01<sup>c</sup> |

#### Effect of pH on bacterial growth

The level of microbial activity in the soil is usually affected by the pH of the soil. In the present study, isolated actinomycetes strains were grown at different pH ranges of 3, 5, 7, 9, and 11 (**Table 6**). The results revealed that optimum growth at OD_600_ for all tested bacterial isolates was observed between the pH of 5 and 9. At pH 3, isolates S4 (0.29 ± 0.11) and S12 (0.25 ± 0.05) had significantly (p < 0.01) higher growth rates compared to other isolates. Isolate S4 was able to withstand the highest pH of 11 having its optimum growth of 1.03 ± 0.07. Growth rate was significantly (p < 0.01) higher in isolate S11 (0.83± 0.15) at pH 5, with the least growth rates recorded for S20 (0.26 ± 0.08), R15 (0.29 ± 0.10) and R11 (0.33 ± 0.09) respectively. At pH 7 and 9, isolate S12 had the highest growth rates (1.19 ± 0.39) and (1.20 ± 0.32), respectively. There were no significant differences in the growth rate of other isolates at pH 9 (**Table 6**). Only 42.85% of bacterial isolates were able to grow at the highest pH of 11(S4 (1.03 ± 0.07), S7 (0.98 ± 0.26), and isolate S20 (0.94 ± 0.17).

**Table 6.**
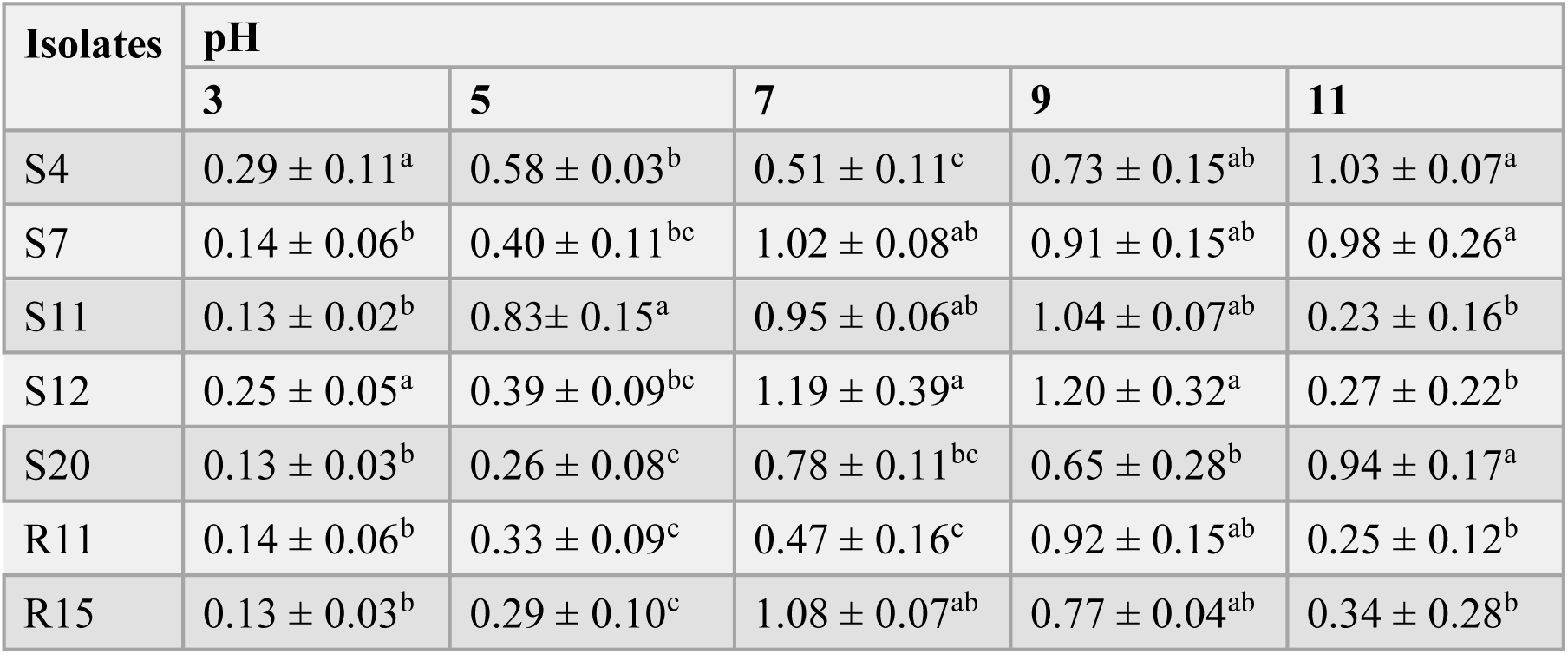
Effect of various pH on the growth of bacterial isolates.

| Isolates | pH |  |  |  |  |
| --- | --- | --- | --- | --- | --- |
|  | 3 | 5 | 7 | 9 | 11 |
| S4 | $0.29 \pm 0.11^a$ | $0.58 \pm 0.03^b$ | $0.51 \pm 0.11^c$ | $0.73 \pm 0.15^{ab}$ | $1.03 \pm 0.07^a$ |
| S7 | $0.14 \pm 0.06^b$ | $0.40 \pm 0.11^{bc}$ | $1.02 \pm 0.08^{ab}$ | $0.91 \pm 0.15^{ab}$ | $0.98 \pm 0.26^a$ |
| S11 | $0.13 \pm 0.02^b$ | $0.83 \pm 0.15^a$ | $0.95 \pm 0.06^{ab}$ | $1.04 \pm 0.07^{ab}$ | $0.23 \pm 0.16^b$ |
| S12 | $0.25 \pm 0.05^a$ | $0.39 \pm 0.09^{bc}$ | $1.19 \pm 0.39^a$ | $1.20 \pm 0.32^a$ | $0.27 \pm 0.22^b$ |
| S20 | $0.13 \pm 0.03^b$ | $0.26 \pm 0.08^c$ | $0.78 \pm 0.11^{bc}$ | $0.65 \pm 0.28^b$ | $0.94 \pm 0.17^a$ |
| R11 | $0.14 \pm 0.06^b$ | $0.33 \pm 0.09^c$ | $0.47 \pm 0.16^c$ | $0.92 \pm 0.15^{ab}$ | $0.25 \pm 0.12^b$ |
| R15 | $0.13 \pm 0.03^b$ | $0.29 \pm 0.10^c$ | $1.08 \pm 0.07^{ab}$ | $0.77 \pm 0.04^{ab}$ | $0.34 \pm 0.28^b$ |

#### Effect of sodium chloride (NaCl) on bacterial growth

Bacterial isolates S4, S7, R11, S11, S12, R15, and S20 were assessed for their ability to withstand salinity stress by growing at different concentrations of sodium chloride (NaCl) 0, 0.2, 0.4, 0.6, 0.8, 1.0 (**Table 7**). Significant differences were observed in the growth of bacterial isolates at the different concentrations of NaCl except at 0.8% NaCl. At 0.2% NaCl, isolates S4 (0.83 ± 0.13) and S20 (0.81 ± 0.10) had their optimum growth which was significantly higher than other isolates evaluated. Likewise, at 0.3% NaCl, optimum growth was recorded in isolates S11 (0.97 ± 0.07), S12 (1.11 ± 0.30), and R15 (1.01 ± 0.02) (**Table 7**). The growth of isolate S7 (1.13 ± 0.13) was significantly highest at 0.4% NaCl than the other isolates, which had low growth. At the highest concentration of 1% NaCl, S7 also stood out as the isolate with the highest growth (0.81 ± 0.04), which could be an indication of its halotolerant nature.

**Table 7.** Effect of NaCl on the growth of bacterial isolates.

| Isolates | NaCl Concentrations (%) |  |  |  |  |  |
| --- | --- | --- | --- | --- | --- | --- |
|  | 0.2 | 0.3 | 0.4 | 0.6 | 0.8 | 1 |
| S4 | $0.83 \pm 0.13^a$ | $0.60 \pm 0.06^{bc}$ | $0.28 \pm 0.15^b$ | $0.53 \pm 0.14^{bc}$ | $0.35 \pm 0.24^c$ | $0.26 \pm 0.14^c$ |
| S7 | $0.34 \pm 0.05^c$ | $0.89 \pm 0.12^{ab}$ | $1.13 \pm 0.13^a$ | $0.36 \pm 0.09^c$ | $0.42 \pm 0.24^c$ | $0.81 \pm 0.04^a$ |
| S11 | $0.56 \pm 0.15^b$ | $0.97 \pm 0.07^a$ | $0.49 \pm 0.18^b$ | $0.55 \pm 0.15^{bc}$ | $0.56 \pm 0.16^b$ | $0.52 \pm 0.17^b$ |
| S12 | $0.43 \pm 0.05^c$ | $1.11 \pm 0.30^a$ | $0.56 \pm 0.17^b$ | $0.52 \pm 0.11^{bc}$ | $0.79 \pm 0.03^a$ | $0.59 \pm 0.13^{ab}$ |
| S20 | $0.81 \pm 0.10^a$ | $0.53 \pm 0.08^c$ | $0.28 \pm 0.21^c$ | $0.69 \pm 0.14^b$ | $0.56 \pm 0.12^b$ | $0.54 \pm 0.07^b$ |
| R11 | $0.33 \pm 0.15^c$ | $0.46 \pm 0.18^c$ | $0.23 \pm 0.16^c$ | $0.42 \pm 0.14^c$ | $0.59 \pm 0.37^b$ | $0.53 \pm 0.26^b$ |
| R15 | $0.38 \pm 0.15^c$ | $1.01 \pm 0.02^a$ | $0.49 \pm 0.20^b$ | $1.00 \pm 0.08^a$ | $0.60 \pm 0.38^b$ | $0.51 \pm 0.19^b$ |

#### Drought tolerance abilities of bacterial isolates

The potentials of the selected bacterial isolates to tolerate drought was evaluated based on concentration and time. The level of tolerance to various concentrations of PEG 8000 by each bacterial isolate was determined as a function of time [35], after inoculation on PEG containing medium. Growth varied among the isolates and depended mainly on the concentration of PEG, as bacterial growth decreased in PEG supplemented medium for all the isolates, irrespective of the concentration, compared to their controls. Data obtained from the growth of most bacterial isolates was higher at lower concentrations than at higher concentrations.

It was also observed that time influenced the tolerance capacities of these isolates as better growth on PEG medium was observed with an increase in time (**Fig 3A-G**). The result obtained for each concentration was compared with that of the control (without PEG). Isolate S12 had maximum tolerance to PEG treatment at 5% PEG (1.53 ± 0.18) after 120 h of growth, with the lowest growth recorded at 20% PEG after 24 h (**Fig 3A**). However, for isolate R11, maximum tolerance was obtained at 5% PEG at 96 h (1.45 ± 0.16), while the lowest tolerance to PEG treatment was recorded in medium containing 20% PEG (0.07 ± 0.33), at 48 h (**Fig 3B**). For isolate S11, growth increased steadily at 15% PEG from 72 h to 120 h, showing tolerance of the isolate to 15% PEG with its optimum growth reached (**Fig 3C**). Isolate R15 achieved maximum tolerance to 5% PEG at 96 h (1.53 ± 0.18). Also, at 96 h, the level of tolerance to 5% PEG and 10% PEG were steady at the same rate. However, it had its lowest growth at 20% PEG at 24 h. Tolerance to 20% PEG steadily increased from 48 h after culture to 120 h. Isolate R15 was able to withstand 5%, 10%, and 20% PEG at 120 h. (**Fig 3D**). The maximum growth (1.07 ± 0.02) on the PEG medium for isolate S4 was reached at 5% PEG at 96 h, while the lowest (0.24 ± 0.23) was reached at 20% PEG at 24 h. However, at 120 h, the isolate was able to tolerate 15% PEG at 20 h (**Fig 3E**). Maximum tolerance for isolate S20 (1.45 ± 0.16) was achieved at 5% PEG concentration at 48 h and remained steady to 120 h, whereas the lowest growth (0.10 ± 0.32) was observed at 20% PEG at 24 h (**Fig 3F**). In the case of isolate S7, its highest tolerance (1.20 ± 0.07) to water stress was also achieved at 5% PEG at 120 h, while the lowest tolerance (0.07 ± 0.33) was reached at 20% concentration at 24 h (**Fig 3G).**

**Fig 3.**
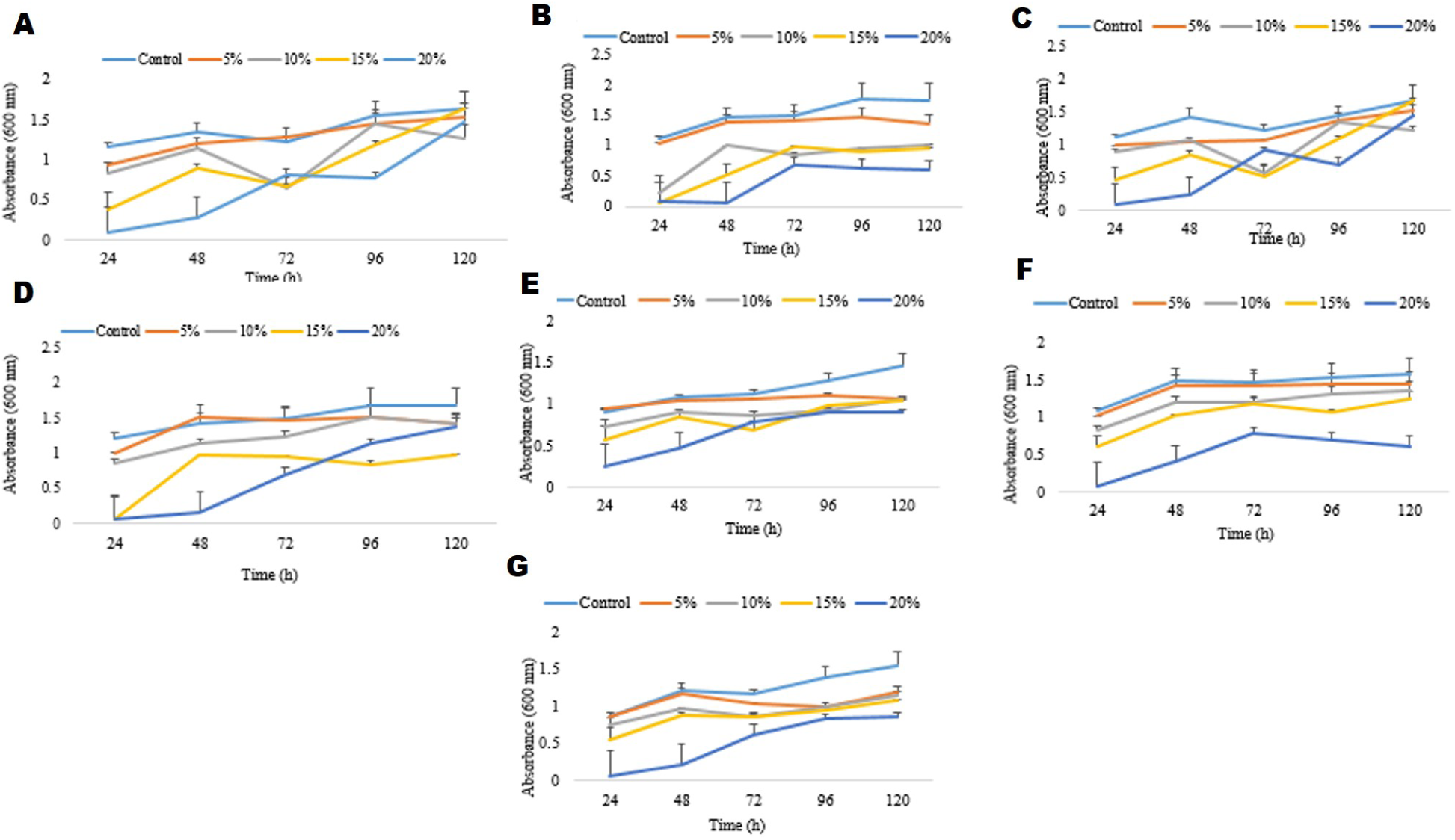
Growth of bacterial isolates (A) S12, (B) R11, (C) S11, (D) R15, (E) S4, (F) S20, and (G) S7 at a different time interval and PEG concentrations

#### Seed germination assay

The bacterial isolates S20, S12, R11, and S11 were tested for their ability to enhance maize growth under drought stress by growing on PEG 8000 medium (−0.30 MPa). Results showed that inoculated seedlings performed better in terms of an increase in growth parameters (shoot and root lengths) than the control (**Fig 4A**). Highly significant differences (p < 0.001) were observed in the root lengths of the various treatments. The watered maize seeds (10 ml sterile distilled water) inoculated with isolate S20 (S20+H_2_0) had significantly longer root lengths than the other seeds inoculated with other isolates and the un-inoculated seeds treated with water. The root length increased by 114.75% more than the un-inoculated watered maize seeds (M+H_2_0) and by 40.37%, 13.26%, and 14.58% more than the root lengths of watered maize seeds inoculated with isolates S11, S12 and R11, respectively. Likewise, isolates S12 and S20 treated with PEG 8000 (−0.30 MPa) also produced longer root lengths than the PEG treated maize seeds inoculated with other isolates and the un-inoculated PEG treated maize seeds (**Fig 4A**). Root length was increased by 110.53% in PEG treated maize seeds inoculated with isolate S20 (S20+PEG) than the un-inoculated PEG treated maize seeds (M+PEG). PEG treated maize seeds inoculated with isolate S12 (S12+PEG) also had long root length which was just 11.11% shorter than the root length of seeds inoculated with isolate S20, but 86.53% and 65.89% longer than the roots of PEG treated seeds inoculated with isolates S11 and R11 respectively. Nevertheless, no significant difference was recorded in the root length of maize seeds inoculated with isolate S11 (1.93±0.19 cm) and that of the un-inoculated maize seeds (1.90 ± 0.15 cm). Highly significant (p < 0.001) differences were also observed in the shoot lengths of all inoculated maize seeds treated with PEG compared to the un-inoculated seeds (**Fig 4A**). Shoot length was significantly highest in watered maize seeds, and PEG treated maize seeds inoculated with isolate S20 than the other treatments. Shoot length increased by 44.82% and 115.61% than the un-inoculated watered maize seeds, and PEG treated maize seeds, respectively. Generally, the root and shoot lengths were shorter in the un-inoculated watered, and PEG treated maize seeds.

**Fig 4.**
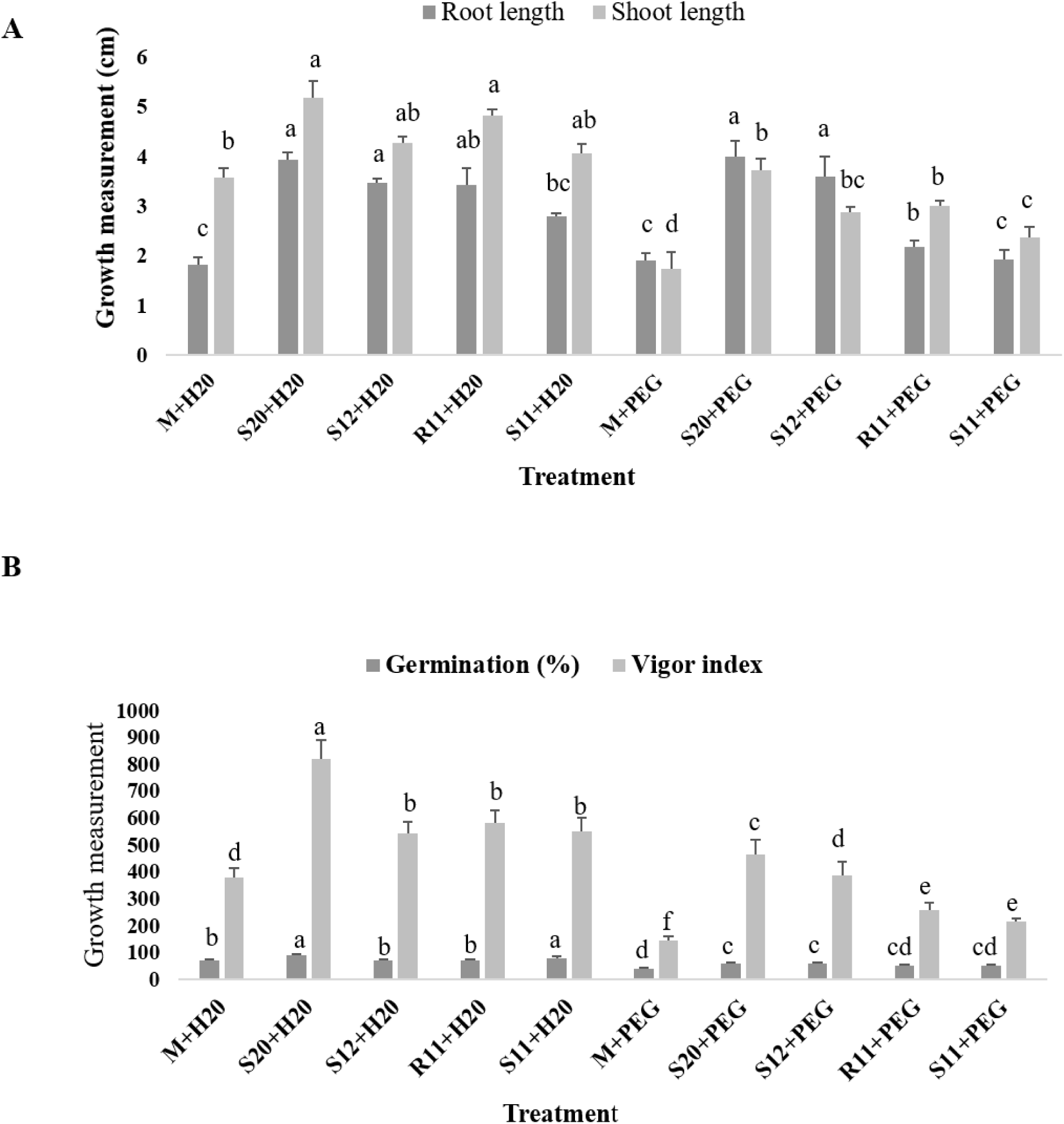
(a) Root and shoot lengths of inoculated and un-inoculated maize seeds treated with either water or PEG 8000, (b) Germination percentage and vigor index of inoculated and un-inoculated maize seeds treated with either water or PEG 8000. M = maize seeds, H_2_0 = water, S20, S12, R11, S11 = maize seeds inoculated with bacterial isolates, and PEG = PEG treated seeds either inoculated or un-inoculated. All values are means of triplicate determinations ± S.E. Means followed by the same letters are not significantly different at P ≤ 0.05 according to New Duncan’s Multiple Range Test (DMRT).

Regarding germination percentage and vigor index, highly significant (p < 0.001), differences were obtained among the treatments (**Fig 4B**). The germination percentage of the PEG treated maize seeds inoculated with isolate S20 was significantly highest than other PEG treated seeds inoculated with isolates S11, S12, and R11 and the control. The germination percentage was higher than the un-inoculated PEG treated maize seeds by 37.53%. The same trend was observed in the watered maize seeds inoculated with isolate S20, which had significantly higher germination percentage than the watered seeds inoculated with isolates S11, S12, and R11. Germination percentage in the S20 inoculated watered seeds was 23.86% higher than the un-inoculated watered seeds (M+H20). Vigor index in the watered treatments was significantly highest in maize seeds inoculated with isolate S20 (S20+H_2_0). This was 108.71% higher than the control seeds (M+H_2_0). Maize seeds inoculated with the same isolate (S20) at water stress (−0.30 MPa), had increased vigor index of 194.81% than the un-inoculated PEG treated seeds (control). All the seeds inoculated with the various bacterial isolates had a significantly higher vigor index than the un-inoculated maize seeds treated with sterile water and PEG medium (**Fig 4B**).

## Discussion

Based on the morphological, biochemical, and 16S rRNA BLAST query of the seven actinomycetes isolates, it was established that 71.43% of the actinomycetes strains isolated from maize rhizosphere soil belong to the genus *Streptomyces*, 14.29% from *Arthrobacter* and 14.29% from the genus *Microbacterium.* The molecular characterization of the isolates was in line with the results obtained from the biochemical tests performed on the various isolates.

The tested bacterial isolates were compared with the description of the cultural, biochemical, and physiological properties of other actinomycetes that fall on the same genus, hence the conclusion of the group [29, 32]. Similar studies have confirmed the dominance of *Streptomyces* spp. isolated from different hosts [45]. Akond, Jahan [46] isolated actinomycetes strains *Streptomyces* and *Nocardia* spp. from straw and compost samples of Savar in Bangladesh. Sreevidya, Gopalakrishnan [47] also isolated actinomycetes from vemicompost and chickpea rhizospheric soils for yield improvement in chickpea.

The phylogenetic analysis of the drought-tolerant actinomycetes isolates used in the present study revealed that these isolates expressed a very high similarity value (99-100%), which was above the 70% borderline degree of relatedness or identity as suggested by Wayne, Brenner (48). Furthermore, the similarities expressed by these isolates with the reference taxa belonging to different species is because of the high similarity value exhibited in the values of DNA re-association, which was above the 70% threshold values [37, 49]. According to Konstantinidis and Stackebrandt (50), this shows a very high genetic similarity that is progressively more consistent, as they cannot be erased overnight [37]. Overall, the high-level branching in the phylogenetic tree is in harmony with the traditional systematic divisions that classifies organisms belonging to the same family and genus into different species [51].

The molecular identification of bacterial isolates is very important because it identifies organisms up to the species level and provides information about the organisms, such as the compounds they produce and whether they are novel or not [52]. The analysis of 16S rRNA gene sequences is a significant method of phylogenetically characterizing microorganisms because it gives a lucid explanation of the evolutionary association between organisms [52, 53]. Most of the closest strains to the bacterial isolates in the present study have been associated with one stress tolerance or the other. For instance, in the study of Srivastava, Patel [54] *Streptomyces rochei* alleviated the stresses caused by salt and *Sclerotinia sclerotiorum* in chickpea. Kanini, Katsifas [55] also reported the antagonistic activity of *S. pseudovenezuelae* against R*hizoctonia solani*. Furthermore, Aly, El Sayed [56] observed the PGP effect of *Streptomyces* spp. on wheat plant under saline conditions. Tolerance to water stress (−0.05 to −0.73 MPa) has also been reported in *Streptomyces geysiriensis* and *Streptomyces coelicolor* which greatly increased the growth and yield of wheat plants subjected to drought stress [54].

An increase in temperature leads to an increase in the activity of cellular enzymes. However, extremely high temperatures cause denaturation of protein structures, while very low temperatures (closer to freezing point) cause the inactivation of enzymes and decrease in cellular metabolism [46]. Most actinomycetes thrive better at temperature ranges between 25 and 30°C. However, certain bacteria require higher temperatures for growth and survival. Examples are the pathogenic bacteria which require 37°C for normal growth and the thermophiles that need a temperature range between 50 to 60°C or more to survive and grow [46]. In this study, a significant decrease in growth was observed for all isolates at 40°C. Nevertheless, 71.43% of the isolates got to their optimum growth at 35°C (S4, S7, S12, R11 and R15). Most actinomycetes species are aerobic, indicating that oxygen is required for their growth. The effect of temperature on the growth of the tested bacterial strains showed that these bacteria could survive at varying temperature ranges. It indicates their potential to survive in temperate, harsh, or hot climate conditions. The results obtained are in agreement with a study by Ndeddy Aka and Babalola [35]. They also reported a decrease in the growth of bacterial isolates when the temperature was raised to 40°C. According to them, the decrease in growth at this temperature could be due to a reduction in metabolic activity of the bacterial isolates caused by the high temperature increase. Everest, Cook [58] reported that *Nocardia* strains isolated from South African soil showed growth at the temperatures 30 and 37°C but did not grow at 45°C. Besides, Bhavana, Talluri [59] reported that *Streptomyces carpaticus* obtained from the sea coast Bay of Vishakhapatnam, Bengal produced optimum mycelial growth and antibiotic at 30°C. However, isolate R11(*Streptomyces indiaensis*), and S12 (*Streptomyces* sp.) grew more than other isolates at 40°C, which may be due to their ability to withstand high temperature, hence may be regarded as high-temperature-tolerant strains.

The pH of an environment influences bacterial survival and growth. For most soil bacteria, the specific pH range is usually between 4 and 9, with the optimum being 6.5 to 7.5 [46], although different bacteria have different pH ranges that favor their growth. Isolates evaluated in this study could adapt and survive at a wide range of pH, which suggests their suitability for drought tolerance enhancement at various soil pH. Kim, Seong [63] also reported the maximum growth of new *Streptomyces* species between the pH ranges of 4.3 to 7.3. Likewise, Palanichamy, Hundet (64) recorded the maximum growth of *Streptomyces* spp. isolated from the Chennai coastal region at pH 7.6 to 8.0. Nevertheless, isolates S4 (*Streptomyces werraensis*), S7 (*Streptomyces luteogriseus*), and S20 (*Streptomyces pseudovenezuelae*) had better growth at the highest pH 11 compared to the other isolates. These isolates may be useful in growth promotion of plants in alkaline environments.

In this study, the bacterial isolates were assessed for tolerance to salinity stress by growing in different concentrations of NaCl. The isolates were able to grow at all concentrations of NaCl. This confirms their ability to withstand various salinity stress conditions and suggests their applicability for salt tolerance improvement in plants. It is also an indication of halotolerant nature of the bacteria strains because of their ability to withstand various levels of salinity stress, especially S7 (*Streptomyces luteogriseus*) which had the highest growth rate at 1% NaCl concentration. A study by Hamid, Ariffin [62] showed that three *Streptomyces* spp. from Malaysian soil grew at a preferred NaCl concentration of 3%. Likewise, Krishnan and Sampath Kumar [63] reported that a *Streptomyces grancidicus* strain isolated from an Indian soil had its optimum activity at 1.5% NaCl concentration, which was also more than the NaCl concentration tested in the present study. Small quantities of salts or metallic ions enhance microbial growth, while large concentrations cause inhibitory effects on growth. High salt concentration also affects osmotic pressure and causes protein denaturation, therefore halophilic bacteria possess specific enzymes in their active configuration that are only activated upon exposure to high salt concentrations [49]. The present study concurs with a study by Ripa, Nikkon (64), who reported that a new *Streptomyces* strain recovered from Bangladesh soil in 2009 exhibited tolerance to salinity stress from 0.5% to 3.0%. Considering isolates that grew profusely at different NaCl concentrations especially isolate S7 which grew profusely at 1% NaCl, they may possess genes that give them the ability to synthesize compatible solutes such as proline or other osmolytes to balance the osmotic potential of their cells under highly saline environments, making them salt-tolerant [65, 66].

For bacteria to improve drought tolerance, tolerance to PEG is required because PEG induces osmotic stress in the supplemented medium [67 68]. The ability of bacteria to grow in this medium will determine its ability to survive and grow in environments with low water availability [67, 68]. Results obtained on drought tolerance abilities of bacterial isolates agree with previous studies on the effect of PEG 8000 on bacterial growth, which confirms that all tested bacterial isolates were able to grow on PEG medium. The ability of some bacteria to resist different concentrations of PEG has been reported. A study by Marasco, Rolli [69] revealed that all bacteria used in the study tolerated 10% and 20% PEG. Moreover, Ali, Sandhya [70] reported that nine out of seventeen fluorescent *Pseudomonas* tested grew at a minimum water potential of -0.30 MPa. The result from this study is in agreement with that of Yandigeri, Meena [57], who reported that *S. olivaceus, S. coelicolor* and *S. geysiriensis* showed significant growth from -0.05 to -0.73 MPa of PEG 6000. The tolerance capacities of these isolates to PEG 8000 reveals their potential to tolerate drought, and this indicates their suitability to be used in improving drought tolerance in plants. In our previous study Chukwuneme, Babalola [26], isolates R15 (*Arthrobacter arilaitensis*) and S20 (*Streptomyces pseudovenezuelae*) with high IAA and ACC deaminase activities grew well at 20% PEG 8000 and mitigated the effect of water stress at 50% and 0% field capacities in maize plants when co-inoculated on vermiculite coated maize seeds. This resulted in greater plant biomass and physiological parameters, especially the chlorophyll content which is relevant in yield increase in plants.

The study on maize seedlings exposed to water stress revealed that significant growths were observed in the root and shoot lengths, germination percentage, and vigor index of the seedlings obtained from seeds inoculated with the bacterial isolates S20, S12, R11, and S11 compared to un-inoculated ones. However, the bacterial isolate S20 (*Streptomyces pseudovenezuelae*) showed outstanding performance in all the parameters measured. Isolate R15 (*Arthobacter arilaitensis*) in the present study, tolerated 5%, 10%, and 20% PEG at 120 h of growth and isolate S20 (*Streptomyces pseudovenezuelae*) significantly increased root length by 110.53% in PEG treated maize seeds (−0.30 MPa) than the un-inoculated control. This confirms the suitability of both isolates as bioinoculants for plant growth promotion of maize under water-deficit conditions. Our findings agree with the study of Yandigeri, Meena [57] who demonstrated that wheat seedlings inoculated with *S. coelicolor, S. olivaceus* and *S. geyiriensis* showed higher seedling vigor as compared to the treatment with cell-free extracts. S20 (*Streptomyces pseudovenezuelae*) in this study had the highest germination percentage and vigor index.

In this study, primers were developed and used to amplify specific genes encoding proteins involved in drought tolerance. The properties of the designed primers concur with primer characteristics of the mercegens-specific PCR primers developed by Aremu and Babalola [37]. They also agree with the characteristics of primers as proposed by Innis, Gelfand [43], which yielded excellent results.

Studies on gene expression are important tools used to understand and compare the universal responses of bacteria to abiotic stress. Bacteria have been found to possess certain genes that enhance their tolerance to abiotic stresses. The designed primer-specific PCR amplifications on total genomic DNA fractions used in this study, targeted some genes responsible for drought tolerance and plant growth promotion in bacteria. Amplification of the target product sizes of drought tolerance and PGP genes by some of the isolates could indicate the presence of these genes in their genome or the DNA chromosomes usually present in the nucleotides of various bacterial cells. Our findings are in harmony with the study of Raddadi, Cherif [71] who reported positive results for all bacterial strains tested for the *accd* gene coding for ACC deaminase enzyme. Ali, Sandhya [70] also reported that isolate SorgP4 amplified the *accd* gene. Bacteria possessing ACC deaminase gene can aid in lessening the ethylene levels in plants and also assist in the protection of plants against certain environmental stresses such as drought, flooding, phytopathogens, and heavy metals [8, 72-74] which induce the synthesis of ethylene. The presence of the a*ccd* gene in all isolates in this study could be of great importance in field applications under stress conditions. The high frequency of occurrence of the *accd* gene in the actinomycetes isolates could also be due to the enzymes ability to catalyze the deamination of other substrates as observed in *Pseudomonas putida* UW4 [75, 76] and *Pyrococcus horikoshii* [76, 77]. The presence of siderophore gene in bacterial isolates can enhance biocontrol of phytopathogenic fungi due to the competition for iron by plants, and also improve the availability of iron to plants. Masalha, Kosegarten [78] reported the role of soil microorganisms in the acquisition of iron and plant growth promotion. All the isolates assessed in this study possessed one or more drought-tolerant and PGP gene. This indicates their potential for possible use as bio-inoculants, not only to improve drought tolerance in plants but also to serve as biofertilizers and biocontrol agents to facilitate plant growth.

The amplification of specific genes encoding proteins involved in drought tolerance by the primers: *GPX, DSP, GTP*, and *GRP* was performed from genomic DNA template fractions from all bacterial strains. However, no sequencing analysis was performed to match them with available genes encoding drought tolerance in GenBank database, hence the need for further investigation. Also, the utilization of novel technologies like the real time-polymerase chain reaction (RT-PCR) could give better results to quantify the levels at which these genes were expressed. The products obtained from the various gene expressions could also be used in the engineering of new drought-tolerant bacteria as well as in the construction of biosensors.

## Acknowledgments

Chukwuneme C.F. acknowledges financial support from the National Research Foundation, South Africa (Grant Nos.UID99457, UID107778). Babalola O. O. acknowledges research support from the National Research Foundation, South Africa (Grant No. UID1236340).

## Author contribution statement

CFC, OOB, FRK, and OBO, contributed to the data collection, wet laboratory, analyses, and drafting of the manuscript for publication. All authors reviewed and approved the final manuscript.

## Data availability statement

All data generated or analyzed during this study are included in this manuscript and its supplementary information file.

## SUPPLEMENTARY INFORMATION

**Table S1:**
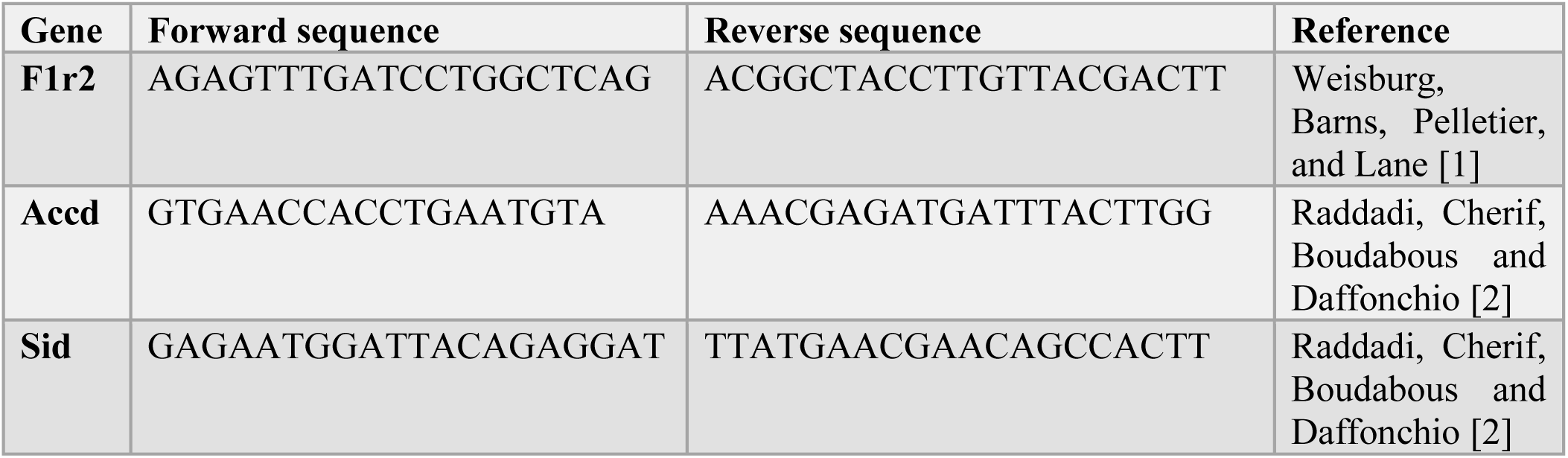
Oligonucleotide primers for PCR amplification of 16S and PGP genes.

**Table S2.**
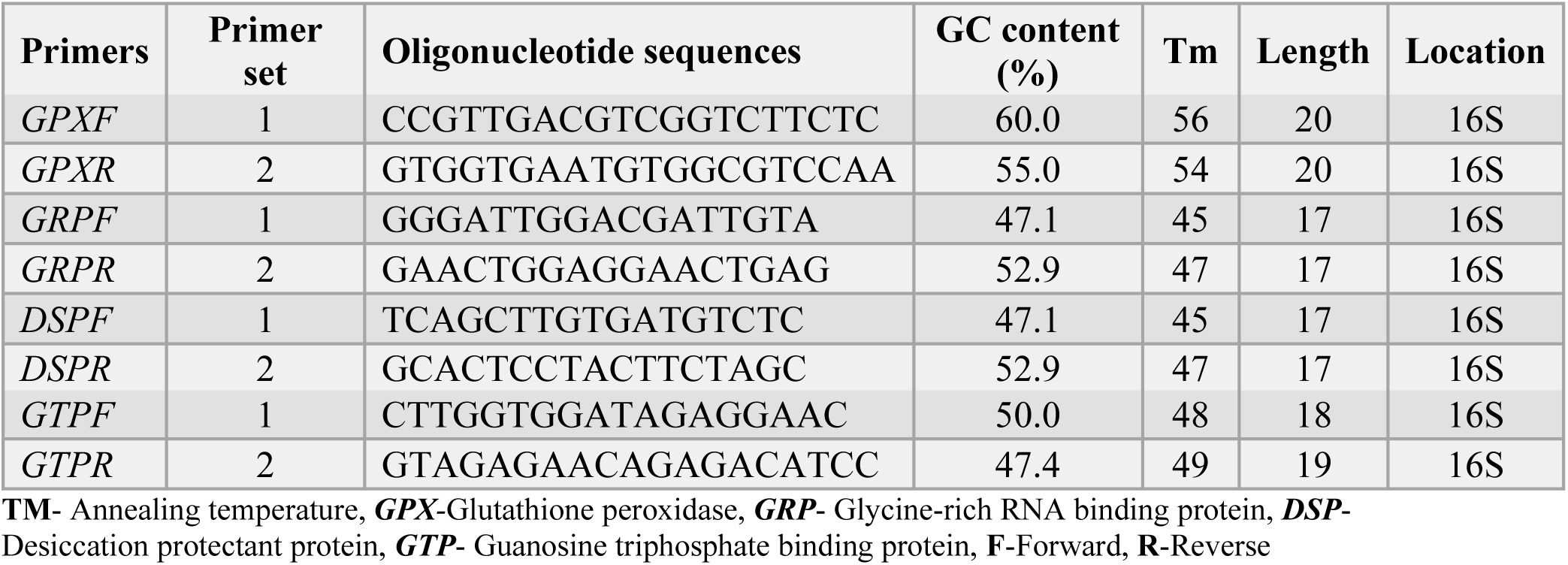
Properties of designed primers for PCR.

**Fig S1.**
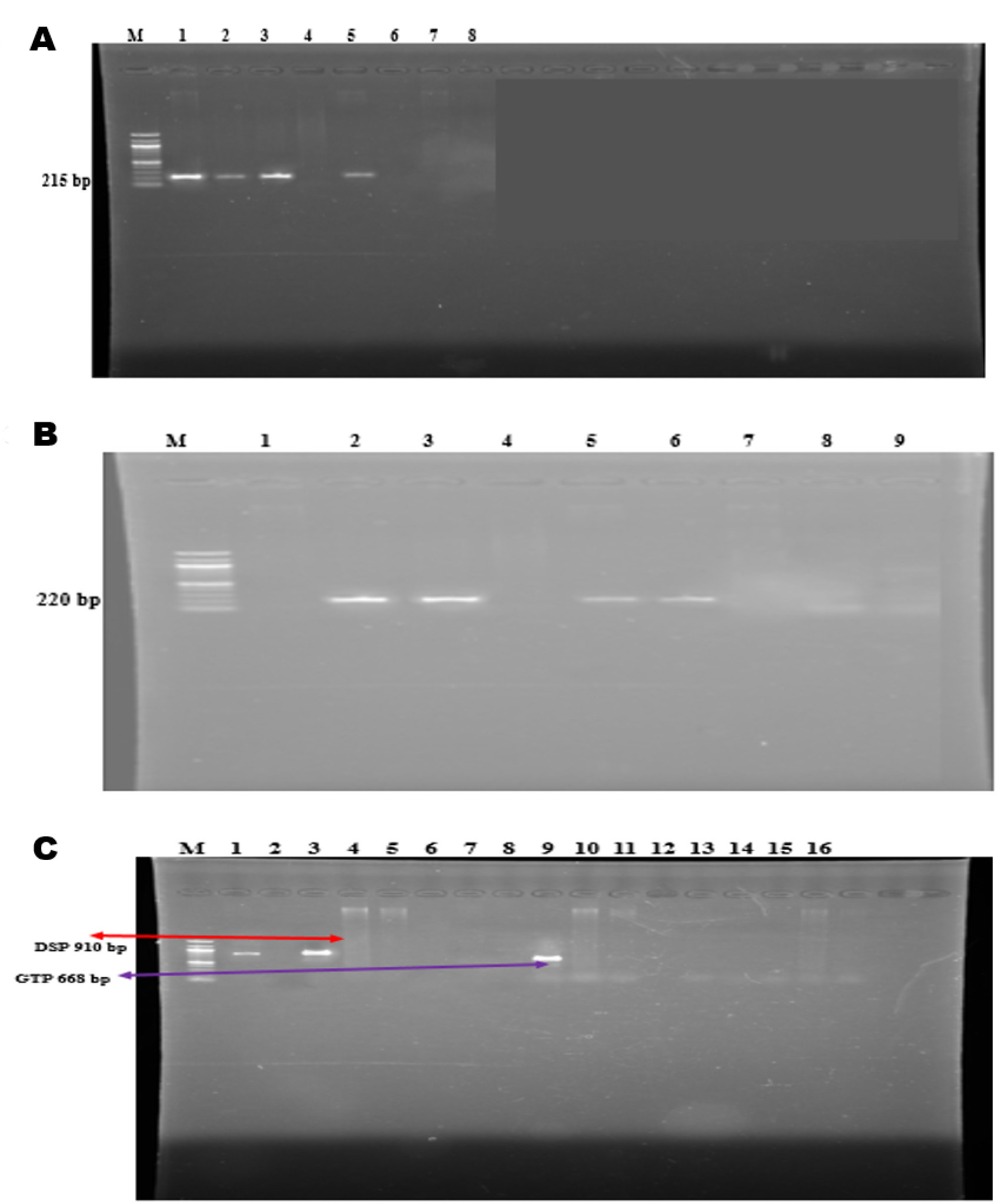
Full length agarose gel electrophoresis of PCR products of designed primers encoding drought tolerance genes. **Legend:** (A) ***GPX*.** Lane M = 100 bp DNA ladder (MWL), Lane 1: R15, Lane 2: S20, Lane 3: S7, Lane 4: S11, Lane 5: R11, Lane 6: S12, Lane 7: S4, (B) ***GRP***. Lane M = 100 bp MWL, Lane 1: S7, Lane 2: R11, Lane 3: S11, Lane 4: S12, Lane 5: S20, Lane 6: R15, Lane 7: R11; (C) ***DSP*** and ***GTP***. For DP, lane M = 100 bp molecular ladder, Lane 1: S20, Lane 2: S11, Lane 3: R15, Lane 4: S4, Lane 5: S7, Lane 6: S12, Lane 7: R11. For GTP, Lane 9 = R15, Lane 10 =S7, Lane 11 = S11, Lane 12 = S12, Lane 13 = S20, Lane 14 = S4 and Lane 15 = R11.

**Fig S2.**
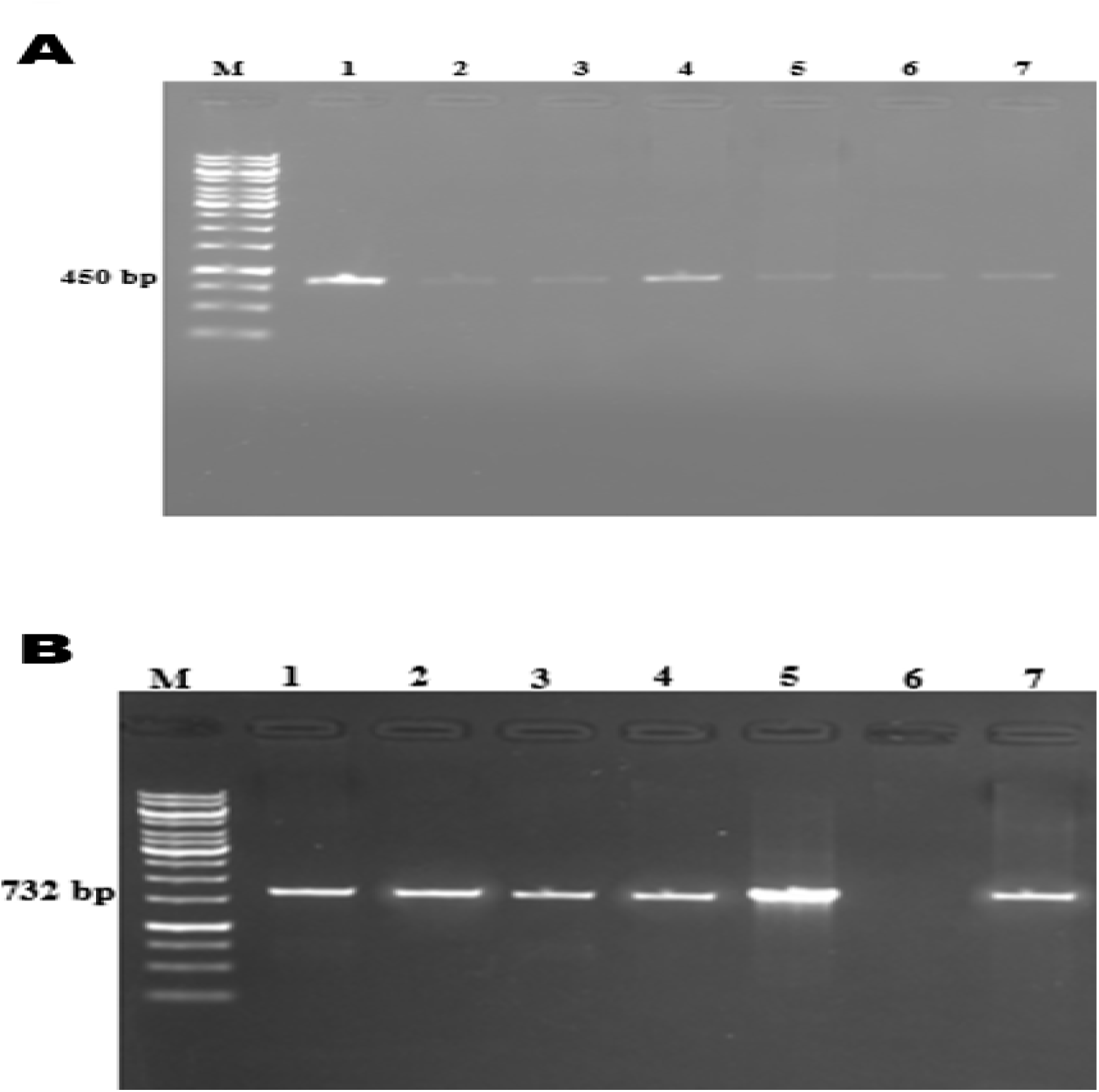
PCR amplification of PGP genes for. **(A)** ACC deaminase (*accd* gene) Lane M = 1 Kb MWL, Lane 1: S11, Lane 2: S12, Lane 3: S7, Lane 4: S20, Lane 5: S4, Lane 6: R15, and Lane 7: R11; **(B)** Siderophore (*Sid* gene) Lane M= 1Kb MWL Lane 1: S20, Lane 2: R11, Lane 3: S12, Lane 4: R15, Lane 5: S11, Lane 6: S7, Lane 7: S4

